# Neuronal selectivity and geometric alignment in the human hippocampus support abstract generalization

**DOI:** 10.64898/2026.08.25.746980

**Authors:** Armin Hakkak Moghadam Torbati, Narges Davoudi

## Abstract

Abstract representations allow the brain to extract shared structure across different experiences and generalize knowledge beyond individual situations. Although previous studies have shown that representational geometry plays a critical role in supporting abstraction, it remains unclear how the composition of neuronal populations gives rise to such generalizable representations. Here, we investigated how neuronal selectivity shapes the emergence of abstract representations by combining a controlled computational model with analyses of human hippocampal single-neuron recordings. We first manipulated the composition of artificial neural populations to test whether increasing task-related information alone is sufficient to improve cross-context generalization. Although increasing stimulus-and response-selective neurons enhanced encoding strength, it did not improve generalization across contexts. In contrast, introducing category-selective neurons increased cross-context generalization, demonstrating that the type of information represented by a population is critical for abstraction. Analyses of human hippocampal neurons revealed a similar principle: category-like and identity-like neurons produced comparable increases in stimulus encoding, but category-like neurons produced substantially stronger improvements in cross-context generalization. Further analyses showed that category-like neurons influenced abstraction by reshaping population geometry. Specifically, category-axis alignment across contexts, rather than the strength of category-related separation, was the geometric property most strongly associated with generalization. Mediation analysis further indicated that category-like neurons contribute to abstraction primarily through their ability to increase geometric alignment across contexts. Together, these findings reveal a population-level mechanism linking neuronal selectivity to abstract computation and suggest that flexible generalization depends not simply on increasing neural information, but on organizing information into geometries that preserve task-relevant relationships across changing conditions.

## Introduction

Consider learning to navigate two different buildings, such as a hospital and a university campus. Although their visual appearance and layout are different, both environments contain common structural features, such as locations connected by routes and places that serve specific purposes. The ability to recognize this shared structure and use it in a new environment without learning everything from scratch reflects the formation of abstract representations and enables generalization across different contexts [1]. Such representations capture task-relevant structure while discarding irrelevant variability in sensory inputs, allowing flexible learning, adaptation, and inference [2–4]. Understanding how the brain constructs abstract representations can therefore provide fundamental insights into the neural computations that support generalization, inference, and flexible behavior [2, 4, 5].

Recent work has shown that abstract representations can be characterized at the population level through representational geometry. In particular, Cross-Condition Generalization Performance (CCGP) provides an operational measure of abstraction by quantifying whether a decoder trained on one set of task conditions can generalize to conditions defined by different combinations of other task variables [6]. Using these approaches, researchers have found that representational geometry constrains the computational regime of neural populations, influencing how neural systems balance abstraction, information specificity, and flexible computation [6–8]. Specifically, representational geometry can regulate the balance between abstraction and memory capacity, with geometries that emphasize shared latent structure supporting generalization, whereas geometries preserving richer stimulus-specific information may favor detailed memory storage [9]. Beyond this trade-off, neural populations were also shown to adopt intermediate representational geometries that preserve high CCGP while maintaining sufficient computational flexibility for more complex computations [10]. From a systems and population perspective, representational geometry captures how different brain regions implement similar task information and behavioral performance through distinct underlying neural organizations. For instance, neural populations across different regions organize identical task information into unique geometries, resulting in varying levels of abstraction and generalization [11]. These geometric variations also reveal distinct computational strategies, even when animals exhibit comparable behavioral performance [12]. Furthermore, mixed-selectivity populations can support disentangled representations in which multiple task variables remain independently decodable through high-dimensional population codes [7]. From a regional perspective, such abstract geometries were shown to emerge during learning in the human hippocampus, where the development of high-CCGP representations directly supported inference and flexible generalization [5]. Despite these advances, it remains unclear which properties of neural populations have the greatest influence on the emergence of abstract representational geometries and how different neuronal subpopulations contribute to the formation of high-CCGP representations.

One possible explanation is that abstract representational geometries emerge simply as a consequence of increasing the amount of task-relevant information encoded within a neural population. Under this hypothesis, increasing the number of neurons encoding task-relevant information, regardless of their functional type, would be expected to enhance CCGP and consequently, promote abstraction. However, an alternative possibility is that abstraction depends not simply on the amount of encoded information, but on the selective contribution of specific neuronal subpopulations. In this view, particular neuronal subpopulations may contribute disproportionately to abstraction, such that changes in these populations, rather than increases in encoded information alone, determine the emergence of high-CCGP representational geometries. Furthermore, both hypotheses predict changes in representational geometry, but it remains unclear whether changes in CCGP are driven primarily by the strength of the category axis or by its alignment across task conditions. Identifying the population-level mechanisms that give rise to abstract representations is essential for understanding how neural circuits support flexible generalization and for uncovering the computational principles that organize representational geometry in biological and artificial neural systems.

To test these hypotheses, we carried out the study in two steps. First, we developed a simple computational toy model that allowed us to systematically manipulate neural populations under controlled conditions. We then tested whether the same principles could explain abstract representations in real hippocampal recordings. Using both the toy model and real hippocampal data, we changed the relative proportions of different neuronal subpopulations and examined how these changes influenced representational geometry, information encoding, and CCGP. We then tested whether abstraction was better explained by (i) the alignment of population activity with the category axis across contexts or (ii) the overall strength of category-related encoding independent of geometric alignment. Finally, we addressed a key mechanistic question: does increasing the number of task-relevant neurons directly improve abstraction, or does it first reorganize the neural representation in a way that enables cross-condition generalization?

## 2. Method

We conducted this study in two sequential phases. First, we developed computational toy models to establish different conditions and test hypotheses regarding the emergence of abstract representations. We then examined whether the same principles generalize to biological neural populations using real hippocampal recordings previously published in [5].

### 2.1. Toy model analysis

We developed computational toy models to systematically examine how changes in neural population composition influence representational geometry, information encoding, and CCGP. The toy model was designed to match the task structure of the hippocampal dataset in [5], allowing the mechanisms identified under controlled conditions to be directly compared with real neural population recordings.

#### 2.1.1. Task design

The task consisted of four stimuli (A, B, C, and D) presented under two contexts (C1 and C2). Context determined the stimulus-response mapping such that the same stimulus required different responses across contexts. In Context 1, stimuli A and C were associated with a left response, whereas stimuli B and D were associated with a right response. In Context 2, this mapping was reversed, with stimuli A and C associated with a right response and stimuli B and D associated with a left response.

This structure defined an abstract stimulus-pair in which stimuli A and C belonged to one category and stimuli B and D belonged to another category (AC vs BD). Importantly, this category was not defined by individual stimulus identity alone, but emerged from the shared response rule across contexts. This task structure therefore allowed us to assess whether population activity represented the abstract AC-vs-BD stimulus-pair in a form that generalized across task conditions.

#### 2.1.2. Neural population model

To investigate how neuronal selectivity shapes abstract representational geometries under controlled conditions, we constructed a computational neural population model in which neuronal activity was generated as a weighted combination of task variables.

Each neuron received input from all task variables, including stimulus, response, context, and noise, with different neurons exhibiting different dominant selectivity profiles through structured weight distributions.

Neuronal activity was defined as:

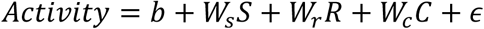

where *S*, *R*, and *C*denote stimulus, response, and context variables coded as ±1, *b* is a constant baseline term, and *ε* ∼ N(0, *σ*^2^)is additive Gaussian noise.

For stimulus-dominant neurons, weights were sampled such that stimulus-related weights were strongly enhanced relative to other variables:

- *W_s_* ∼ N(2.5,0.3)
- *W_r_* ∼ N(0.4,0.2)
- *W_c_* ∼ N(0.4,0.2)

For response-dominant neurons, response weights were dominant:

- *W_s_* ∼ N(0.4,0.2)
- *W_r_* ∼ N(2.5,0.3)
- *W_c_* ∼ N(0.4,0.2)

For context-dominant neurons, context weights were dominant:

- *W_s_* ∼ N(0.4,0.2)
- *W_r_* ∼ N(0.4,0.2)
- *W_c_* ∼ N(2.5,0.3)

For noise-dominant neurons, all task-related weights were close to zero:

- *W_s_* ∼ N(0.05,0.05)
- *W_r_* ∼ N(0.05,0.05)
- *W_c_* ∼ N(0.05,0.05)

This formulation produced a heterogeneous mixed-selectivity population in which all neurons encoded all task variables to varying degrees, while differing in their relative dominance across stimulus, response, and context dimensions.

In addition to this mixed-selectivity population, we introduced a distinct class of category-neurons. These neurons were defined by their tuning to the abstract task structure, responding similarly to stimuli A and C and differently to stimuli B and D, thereby explicitly encoding the AC-versus-BD categorical axis.

#### 2.1.3. Population manipulations

To investigate how changes in neural population composition influence representational geometry and abstraction, we generated three population configurations.

The first configuration represented a context-dominated population and consisted of:

- 1 stimulus-selective neuron
- 1 response-selective neuron
- 60 context-selective neurons
- 38 noise neurons

This configuration was designed to produce a neural population in which task representations were dominated by context information.

In the second configuration, the number of stimulus-and response-selective neurons was increased while maintaining the same number of context-selective neurons. The population consisted of:

- 15 stimulus-selective neurons
- 15 response-selective neurons
- 60 context-selective neurons
- 10 noise neurons

This manipulation allowed us to examine how increasing the numbers of stimulus-and response-selective neurons influenced information encoding, representational geometry, and CCGP.

In the third configuration, we introduced an additional population of category-selective neurons while keeping the stimulus-, response-, and context-selective populations unchanged. The population consisted of:

- 15 stimulus-selective neurons
- 15 response-selective neurons
- 15 category-selective neurons
- 60 context-selective neurons
- 10 noise neurons

This manipulation allowed us to examine how introducing category-selective neurons influenced information encoding, representational geometry, and CCGP.

#### 2.1.4. Population analyses

To characterize abstract representations relevant to the AC vs BD stimulus-pair, we focused our analyses on the geometry of population activity along the category axis.

To quantify abstraction of the AC versus BD categorical structure, we measured CCGP using a linear decoding approach. Specifically, we trained a linear SVM classifier on neural population activity from one context and tested its ability to decode the same variable in another context. CCGP was defined as the classification accuracy of the decoder when trained and tested across different task contexts. To ensure symmetry, decoding was performed in both directions (e.g., Left→Right and Right→Left for response-based splits, or A→B and B→A for stimulus-based splits), and the final CCGP value was computed as the average accuracy across both directions. This procedure was applied separately for each population configuration to assess how changes in neural population composition affect the generalization of AC versus BD representations across task contexts.

To quantify how strongly neural populations encoded task variables, we performed a linear encoding analysis relating neural activity to the task design. This analysis was used to assess whether changes in neural population composition affected the amount of task-related information encoded about stimulus, response, and context, and to compare these changes with the corresponding alterations in representational geometry and CCGP. Neural activity was modeled using a linear regression in which stimulus identity, response, and context were used as predictors. Stimulus identity was represented using a set of four binary predictors corresponding to stimuli A, B, C, and D, while response and context were each represented by a single predictor. All predictors were z-scored prior to model fitting. Encoding strength was quantified at the single-neuron level using regression coefficients and the coefficient of determination (R²). For stimulus encoding, we computed the mean absolute regression coefficient across the four stimulus predictors. Response and context encoding were defined as the absolute regression coefficients of their respective predictors. R² was used as a measure of overall variance explained by the model. Finally, population-level encoding measures were obtained by averaging neuron-wise stimulus, response, and context coefficients, as well as R², across all neurons within each population configuration.

#### 2.1.5. Category neuron sweep

To investigate how category-selective neurons contribute to the emergence of abstract representations, we performed a parametric sweep over the number of category-neurons in the population while keeping all other neuronal subpopulations fixed. Specifically, the number of category-neurons was systematically varied (0, 5, 10, 15, 20, 30, 40, and 60 neurons), while the numbers of stimulus-, response-, context-, and noise-selective neurons were held constant across all simulations.

For each configuration, we re-generated the neural population and repeated the full analysis pipeline, including (i) linear encoding analysis and (ii) CCGP for the AC vs BD stimulus-pair. Encoding strength was quantified using the average absolute regression coefficient (category β), while abstraction was quantified using AC vs BD CCGP computed via cross-condition decoding accuracy.

This procedure allowed us to directly assess how increasing the proportion of category-neurons affects both the strength of encoding and the emergence of abstract, cross-context generalization.

### 2.2. Real hippocampal data

To examine whether the principles identified in the toy model extend to biological neural populations, we analyzed previously published human hippocampal single-unit recordings collected during an inferential reversal-learning task [5]. The dataset was obtained from patients with pharmacoresistant epilepsy undergoing intracranial monitoring with depth electrodes. Participants performed a task requiring them to infer latent changes in task structure from feedback and generalize learned stimulus–response relationships across changing contexts.

The original dataset included recordings from multiple brain regions; however, here we focused specifically on hippocampal neurons because the hippocampus exhibited abstract and disentangled representations of task variables in the original study [5]. Our analysis was designed to test a specific mechanistic hypothesis derived from the toy model: whether abstraction measured by stimulus-pair cross-condition generalization performance (CCGP) is preferentially supported by neurons whose activity is aligned with the abstract category structure, rather than by neurons encoding individual stimulus identity alone.

#### 2.2.1 Dataset and preprocessing

The behavioural paradigm consisted of a serial reversal-learning task in which participants learned stimulus–response–outcome associations under two latent contexts. Each recording session contained four visual stimuli (A–D), with each stimulus associated with a specific response and reward outcome according to one of two possible stimulus–response maps. The active mapping changed unpredictably during the session, requiring participants to infer context switches from feedback and update their behavioural strategy. Importantly, the two contexts were systematically related: stimulus–response associations were inverted between contexts, creating a structured task space in which stimulus identity, context, response, and outcome could be dissociated. Once participants inferred the current context, they could generalize the learned rule to stimuli that had not yet been encountered following the context transition.

For our analysis, we focused on the eight experimental conditions defined by the combination of stimulus identity (four stimuli: A–D) and context (two contexts: C1 and C2). This eight-condition space provides the basis for quantifying stimulus-pair abstraction because stimuli were grouped according to their shared latent structure: stimuli A and C belonged to one abstract category, whereas stimuli B and D belonged to the opposite category (AC versus BD).

Neural activity was quantified from hippocampal single units during the stimulus-processing epoch. In the original dataset, action potentials were counted within defined task epochs, including a stimulus period following stimulus onset, and these firing-rate responses were used for subsequent encoding and population analyses.

#### 2.2.2 Single-neuron encoding and functional classification

To characterize the functional selectivity of individual hippocampal neurons, we first quantified how firing-rate responses were explained by task-related variables. For each hippocampal neuron, trial-by-trial firing rates during the stimulus epoch were modeled using a linear encoding model including stimulus identity, context, response, and reward as predictors:

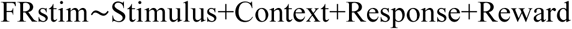

Stimulus identity was treated as a categorical predictor, whereas context, response, and reward were included as additional task variables. For each neuron, the full model was compared with reduced models in which each predictor was removed individually. The reduction in explained variance between the full and reduced models (ΔR²) was used to quantify the unique contribution of each task variable to neuronal activity.

For each predictor, statistical significance was assessed from the encoding model, and neurons were classified according to their significant task-related contributions. Neurons without any significant task-variable modulation were classified as non-selective. When only one task variable significantly explained neuronal activity, neurons were assigned to that functional class. For neurons showing significant modulation by multiple variables, the variable associated with the largest ΔR² was considered the dominant predictor, and neurons were classified according to this dominant contribution. This procedure allowed identification of neurons primarily associated with stimulus, context, response, or reward information while preserving the possibility of mixed selectivity.

#### 2.2.3 Classification of stimulus-related neurons: category-like and identity-like neurons

To determine whether stimulus-related hippocampal neurons preferentially encoded individual stimulus identity or abstract category structure, we characterized their response patterns across the complete stimulus–context space. To avoid circularity between neuronal subtype classification and subsequent population-level analyses, for each neuron, trials were randomly divided into two non-overlapping subsets within each of the eight stimulus–context conditions (four stimuli × two contexts). The first subset was used exclusively for neuronal subtype classification, whereas the second independent subset was reserved for all subsequent analyses. Thus, the trials used to define category-like and identity-like response profiles were never used to evaluate their contribution to population-level generalization.

Using only the classification subset, we characterized the response pattern of each stimulus-related neuron across the complete stimulus–context space. For each neuron, we constructed an eight-dimensional response vector containing the mean firing rate for each stimulus–context combination:

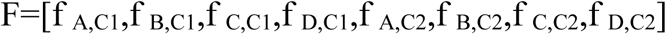

Where each element corresponds to the response of the neuron to one stimulus presented under one of the two task contexts. The response vector was z-scored across the eight conditions before template matching, ensuring that classification was based on the relative organization of responses rather than differences in overall firing magnitude. We quantified similarity between each neuronal response profile and two alternative representational structures: identity-based coding and category-based coding.

Identity coding was assessed using four stimulus-specific templates, one for each stimulus. Each identity template represented a context-invariant preference for a single stimulus, assigning a positive value to the two conditions in which that stimulus appeared across contexts and negative values to all other stimulus conditions. For example, the template for stimulus A was:

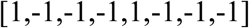

representing a response pattern selective for stimulus A regardless of context.

Category coding was assessed using templates representing the latent category structure of the task. Stimuli A and C belonged to one abstract category, whereas stimuli B and D belonged to the opposite category. Accordingly, the category template assigned the same value to A and C and the opposite value to B and D, while preserving this organization across both contexts:

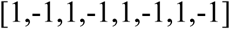

Because the polarity of category preference is arbitrary, the inverse category template was also considered:

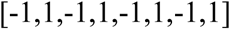

For each neuron, template similarity was quantified using Pearson correlation. The category score was defined as the maximum absolute correlation with the two category templates, whereas the identity score was defined as the maximum absolute correlation across the four identity templates.

Neurons were classified as category-like when their category score exceeded their identity score, indicating that the organization of their responses across the eight conditions was better described by the abstract AC versus BD structure than by a stimulus-specific identity representation. Conversely, neurons were classified as identity-like when their identity score exceeded their category score. Importantly, these subtype assignments were derived exclusively from the classification subset of trials and were subsequently held fixed when the independent analysis subset was used to construct pseudopopulations and quantify cross-condition generalization and population geometry.

#### 2.2.4 Pseudopopulation construction and population manipulations

To test whether the composition of hippocampal neural populations influences abstract cross-context generalization, we constructed pseudopopulations by systematically varying the contribution of identity-like and category-like neurons while maintaining the contribution of other functional neuronal classes.

For each pseudopopulation realization, neurons were randomly sampled from the previously identified functional classes, including identity-like, category-like, context-related, response-related, reward-related, and non-selective populations. Importantly, all population analyses were performed using only the independent analysis subset of trials that was not used for neuronal subtype classification (see Section 2.2.3). The selected neurons were then combined into population response matrices containing trial-level activity across the eight stimulus– context conditions.

In the first manipulation, we defined a baseline population consisting of 13 identity-like neurons together with 56 context-, 13 response-, 13 reward-, and 100 non-selective neurons, while excluding category-like neurons. We then generated controlled population manipulations by adding 5, 10, or 13 category-like neurons.

In the second manipulation, we defined a baseline population consisting of 13 category-like neurons together with 56 context-, 13 response-, 13 reward-, and 100 non-selective neurons, while excluding identity-like neurons. We then generated controlled population manipulations by adding 5, 10, or 13 identity-like neurons.

Across both manipulation schemes, the numbers of context-, response-, reward-, and non-selective neurons were held constant, allowing us to isolate the effect of representational composition on population-level abstraction.

For each population composition, neurons were randomly sampled and combined into pseudopopulations across 100 independent iterations to account for variability arising from neuron selection. For each iteration, the maximum number of trials available per stimulus– context condition (K) was determined from the held-out analysis trials of the selected neurons. Specifically, K was defined as the minimum number of available trials across all eight stimulus–context conditions after restricting analyses to trials that were present for all selected neurons. This procedure ensured balanced sampling across conditions while maximizing the available independent data.

For each iteration, K trials were randomly selected from each stimulus–context condition to construct an 8K × N population activity matrix, where K represents the number of trials per condition and N represents the number of sampled neurons. This approach enabled comparison of population configurations using balanced trial numbers while preserving independence between neuronal classification and population-level analyses.

This experimental design enabled us to test whether improvements in abstraction were attributable to increased stimulus information alone or whether they specifically depended on the inclusion of category-aligned neuronal populations.

#### 2.2.5 Encoding strength and CCGP population-level analyses

After constructing pseudopopulations with controlled variations in neuronal composition, we quantified how these changes affected both the amount of stimulus-related information carried by the selected neuronal populations and their ability to support abstract cross-context generalization.

For each pseudopopulation, stimulus encoding strength was quantified from the single-neuron encoding analysis described above. Specifically, the stimulus-related ΔR² values of the neurons included in each pseudopopulation were summarized to provide a single measure of stimulus information represented by that population. This measure was calculated for each iteration and population composition.

Stimulus encoding was then compared across the identity-like and category-like neuron manipulations to determine whether increasing either neuronal population similarly increased the overall amount of stimulus information.

Abstraction was quantified using CCGP for the stimulus-pair distinction A/C versus B/D. CCGP measures whether the neural representation of this stimulus grouping generalizes across task conditions, with a chance level of 0.5.

Stimulus-pair CCGP was calculated separately for each pseudopopulation and iteration exclusively on held-out analysis trials that were not used for defining neuronal selectivity. CCGP values were averaged across iterations within each pseudopopulation to obtain one CCGP value for each pseudopopulation. We then examined how CCGP changed as increasing numbers of identity-like or category-like neurons were added to the population. By comparing these changes with stimulus encoding strength, we tested whether stronger stimulus encoding was sufficient to improve abstraction, or whether improvements in CCGP depended specifically on the representational type of the added neurons.

#### 2.2.6 Population geometry analysis

##### 2.2.6.1 dPCA-like population geometry

To investigate how neuronal composition shapes the representational geometry underlying stimulus-pair abstraction, we analyzed the organization of population activity across the full eight-condition task space (four stimuli × two contexts). Population geometry was estimated from the held-out analysis subset using the same pseudopopulation compositions used for decoding analyses and 100 random sampling iterations. For each iteration, K trials per stimulus–context condition were randomly sampled, where K was determined as described in Section 2.2.4.

For each iteration, trial-level population activity was organized into an eight-condition response matrix, where each condition corresponded to one stimulus–context combination. The mean response across sampled trials was calculated for each condition, resulting in an eight-condition-by-neuron activity matrix representing the population response structure across the task space. To reduce the influence of differences in firing-rate scale across neurons, responses were z-scored across conditions independently for each neuron.

The condition-level population matrices obtained from the 100 random pseudopopulation realizations were averaged across iterations for each scenario, providing a representative population geometry while preserving the multivariate organization of neuronal responses. PCA was then applied to this averaged population representation to obtain a low-dimensional embedding of the eight task conditions.

The first two principal components were used to visualize the eight stimulus–context conditions (A–D in contexts C1 and C2) in a common low-dimensional population space. Each point represented the mean population response to one stimulus in one context (A-C1, B-C1, C-C1, D-C1, A-C2, B-C2, C-C2, and D-C2), allowing changes in the arrangement of the eight conditions across neuronal-composition scenarios to be directly visualized.

##### 2.2.6.2. Quantification of population geometry

To complement the visualization of population geometry with a quantitative measure, we calculated how strongly the two stimulus groups (AC and BD) were separated in the PC1–PC2 space. This analysis was used to determine whether changing neuronal composition produced a measurable increase in the geometric separation of the two stimulus groups, rather than only a visually apparent change in the PCA plots.

The eight stimulus–context conditions were divided into AC and BD groups. The PC1 and PC2 coordinates of A-C1, C-C1, A-C2, and C-C2 were averaged to obtain a single centroid for the AC group, while the coordinates of B-C1, D-C1, B-C2, and D-C2 were averaged to obtain a single centroid for the BD group. Category separation was then quantified as the Euclidean distance between these two centroids. This procedure yielded one category-separation value for each neuronal-composition scenario, with larger values indicating greater separation of the AC and BD representations in the low-dimensional population space.

#### 2.2.7 Relationship between population geometry and stimulus-pair abstraction

To determine which geometric properties of neural population activity underlie stimulus-pair abstraction, we examined whether changes in CCGP were better explained by the alignment of categorical representations across contexts or by the magnitude of category-related activity. Although previous analyses could determine whether increasing the proportion of category-like neurons enhances stimulus-pair CCGP, they do not reveal the geometric mechanism responsible for this improvement. Therefore, we quantified two complementary properties of the category representation: category-axis alignment, reflecting the consistency of the category structure across contexts, and category-axis strength, reflecting the magnitude of separation between the AC and BD stimulus groups.

For each pseudopopulation realization, category-axis alignment, category-axis strength, and stimulus-pair CCGP were calculated using the same resampling framework described above from held-out population activity (100 iterations per scenario, with K trials per stimulus– context condition). At each iteration, neurons were randomly sampled from identity-like, category-like, context-related, response-related, reward-related, and non-selective populations according to the predefined population composition. The resulting held-out pseudopopulation activity matrix was used to quantify stimulus-pair CCGP and the geometric properties of the categorical representation.

##### 2.2.7.1 Category-axis alignment

Category-axis alignment was quantified to assess whether the neural representation of the AC versus BD stimulus grouping preserved a consistent categorical direction across contexts. This measure captures the extent to which the population-level category structure remains stable when task conditions change, providing a geometric measure of abstraction beyond simple category information strength.

For each pseudopopulation, population activity was organized into the eight stimulus–context conditions (A–D × C1–C2). Within each context separately, category centroids were calculated by averaging the population coordinates of stimuli belonging to the same abstract category. The category axis for each context was defined as the vector separating the AC and BD category centroids:

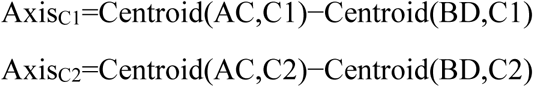

The consistency of the categorical axis across contexts was then quantified using the cosine similarity between the two context-specific category axes:

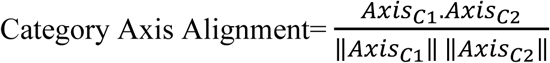

Higher values indicate that the population preserves a more stable categorical organization across contexts.

##### 2.2.7.2 Category-axis strength

To distinguish geometric alignment from the overall magnitude of category-related information, we additionally quantified category-axis strength. This measure captures the separation between AC and BD representations independently within each context.

For each context, the Euclidean distance between the AC and BD category centroids was calculated:

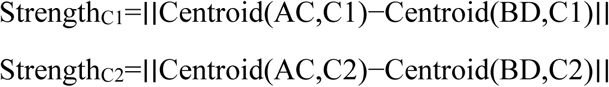

The final category-axis strength was defined as the average separation across the two contexts:

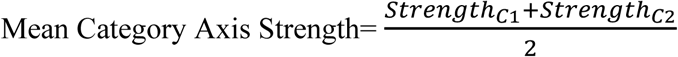

Thus, category-axis strength reflects how strongly the population differentiates AC from BD, whereas category-axis alignment reflects whether this differentiation occurs along a consistent geometric direction across contexts.

##### 2.2.7.3 Statistical relationship with CCGP

To determine which geometric property best explained stimulus-pair abstraction, we evaluated the relationship between stimulus pair CCGP and the two geometric measures.

First, Pearson correlations were calculated between:

1. Category-axis alignment and CCGP
2. Category-axis strength and CCGP

Next, partial correlation analyses were performed to determine whether each geometric measure was independently associated with Category CCGP after accounting for other factors that could influence the relationship. Specifically, the association between category-axis alignment and CCGP was evaluated while controlling for category-axis strength, identity-like neuron number, and category-like neuron number. Conversely, the association between category-axis strength and CCGP was evaluated while controlling for category-axis alignment, identity-like neuron number, and category-like neuron number.

For each partial correlation analysis, the effects of the selected control variables were first removed from both variables of interest using linear regression. Pearson correlation was then calculated between the residuals of the two variables. This approach quantified the unique association between each geometric property and CCGP while accounting for differences in neuronal composition and the alternative geometric feature.

To account for multiple comparisons, false discovery rate (FDR) correction using the Benjamini–Hochberg procedure was applied to the p-values of the partial correlation analyses comparing the independent associations of category-axis alignment and category-axis strength with Category CCGP.

As a sensitivity analysis, we additionally assessed whether the observed zero-order relationships with CCGP were dependent on treating individual pseudopopulation resampling iterations as separate observations. Resampling iterations were therefore aggregated within each neuronal composition, defined by the combination of category-like and identity-like neuron numbers. For each composition, category-axis alignment, category-axis strength, and Category CCGP were averaged across the 100 resampling iterations. Pearson correlations between the composition-level mean CCGP and the mean alignment or mean strength were then calculated. Because only four neuronal compositions were available, this analysis was used specifically to assess the robustness of the zero-order associations to resampling-level dependence, rather than to estimate the independent contributions of alignment and strength. The results of this sensitivity analysis are reported in the Supplementary Results.

To further assess whether the differential associations of category-axis alignment and strength with CCGP were disproportionately influenced by individual category-like neurons, we performed a leave-one-category-neuron-out sensitivity analysis. Each of the 13 category-like neurons was removed in turn, and the pseudopopulation analysis was repeated using the remaining category-like neurons. For each leave-one-out dataset, four neuronal compositions were evaluated (0, 5, 10, or 12 category-like neurons, with the number of identity-like neurons fixed at 13), with 100 pseudopopulation realizations generated per composition using the same held-out trials and analysis procedures as in the primary analysis. Partial correlations between category-axis alignment and CCGP were then recomputed while controlling for category-axis strength and category-like neuron number; conversely, partial correlations between category-axis strength and CCGP were recomputed while controlling for category-axis alignment and category-like neuron number. Identity-like neuron number was not included as a covariate in this sensitivity analysis because it was constant across all compositions. The difference between the alignment–CCGP and strength–CCGP partial correlations was calculated for each leave-one-out dataset. This analysis was used as an influence analysis to determine whether the stronger independent association between category-axis alignment and CCGP was robust to the removal of any single category-like neuron. Results are reported in the Supplementary Results.

#### 2.2.8 Mediation analysis

To determine whether category-like neurons influence stimulus-pair abstraction directly or through changes in population geometry, we performed a mediation analysis testing whether category-axis alignment mediates the relationship between the number of category-like neurons and CCGP.

The motivation for this analysis was that increasing category-like neurons could improve abstraction through two different mechanisms. First, category-like neurons may simply provide additional category-related information, leading to a direct increase in decoding performance. Alternatively, these neurons may reorganize the population representation by creating a more stable category axis across contexts, which subsequently enables better cross-condition generalization. Therefore, we tested whether category-axis alignment represents the intermediate geometric mechanism linking category-like neuronal composition to abstraction.

For each independent pseudopopulation realization, the number of category-like neurons was used as the predictor variable (X), category-axis alignment as the mediator (M), and CCGP as the outcome variable (Y). All variables were obtained from held-out population activity. The number of identity-like neurons and category-axis strength were included as covariates to control for possible effects of population composition and overall category-related signal magnitude.

First, we tested whether the number of category-like neurons predicted category-axis alignment (path a) using a linear regression model:

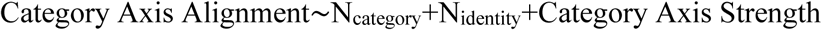

A significant positive relationship would indicate that adding category-like neurons systematically increases the consistency of the category axis across contexts.

Second, we tested whether category-axis alignment predicted CCGP while controlling for category-like neuron number and the covariates (path **b** and direct effect **c′**):

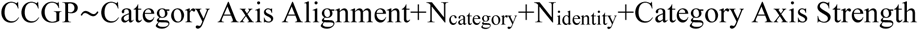

This analysis determined whether category-axis alignment explained additional variance in abstraction beyond the number of category-like neurons alone.

Third, we estimated the total effect (**c**) of category-like neuron number on CCGP using a regression model without the mediator:

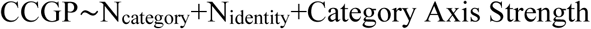

The indirect mediation effect was calculated as the product of path **a** and path **b**:

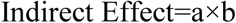

To assess the statistical reliability of the indirect effect, we performed non-parametric bootstrap resampling with 5000 iterations. In each bootstrap iteration, pseudopopulation realizations were randomly resampled with replacement, the mediation paths were recalculated, and the distribution of the resulting indirect effects was used to estimate the 95% confidence interval. The mediation effect was considered significant when this confidence interval did not include zero. This analysis tested whether the influence of category-like neurons on stimulus-pair abstraction is explained by their ability to establish a stable categorical geometry across task contexts.

## 3. Results

### 3.1. Toy-model population manipulations alter task-variable encoding and cross-condition generalization

We first examined how progressive changes in population composition affected task-variable encoding and cross-condition generalization in the toy model. Increasing stimulus and response neurons resulted in an increase in stimulus encoding from β = 0.031 to β = 0.077 and response encoding from β = 0.163 to β = 0.511, while context encoding remained comparable (β = 1.522 vs β = 1.566; Table 1). Mean encoding R² increased from 0.536 to 0.723. Context CCGP remained at 1.000, and response CCGP increased from 0.963 to 1.000, whereas stimulus-pair CCGP remained unchanged at 0.500.

**Table 1.** Summary of encoding strength and CCGP across toy-model population configurations.

| Population configuration | Stimulus beta | Response beta | Context beta | Category beta | Mean encoding $R^2$ | Context CCGP | Stimulus-pair CCGP (AC vs BD) | Response CCGP |
| --- | --- | --- | --- | --- | --- | --- | --- | --- |
| Context-dominated | 0.031 | 0.163 | 1.522 | --- | 0.536 | 1.000 | 0.500 | 0.963 |
| Increased stimulus/response neurons | 0.077 | 0.511 | 1.566 | --- | 0.723 | 1.000 | 0.500 | 1.000 |
| Added category-selective neurons | 0.157 | 0.460 | 1.352 | 0.227 | 0.741 | 1.000 | 0.619 | 1.000 |

The introduction of category-selective neurons produced additional changes in population-level encoding and cross-condition generalization. Category encoding reached β = 0.227, while stimulus encoding further increased from β = 0.077 to β = 0.157. Response encoding remained relatively constant, while context encoding decreased from β = 1.566 to β = 1.352, and mean encoding R² from 0.723 to 0.741. Context and response CCGP remained at 1.000, whereas stimulus-pair CCGP increased from 0.500 to 0.619.

### 3.2. Category-neuron number differentially affects encoding strength and CCGP in the toy model

To further examine how the number of category-selective neurons influences category encoding and CCGP, we systematically varied the number of category-selective neurons while keeping the remaining neuronal populations constant. Increasing the number of category-selective neurons produced a monotonic increase in category encoding coefficient (beta), with values progressively increasing across the tested population sizes (Figure 1A).

**Figure 1.**
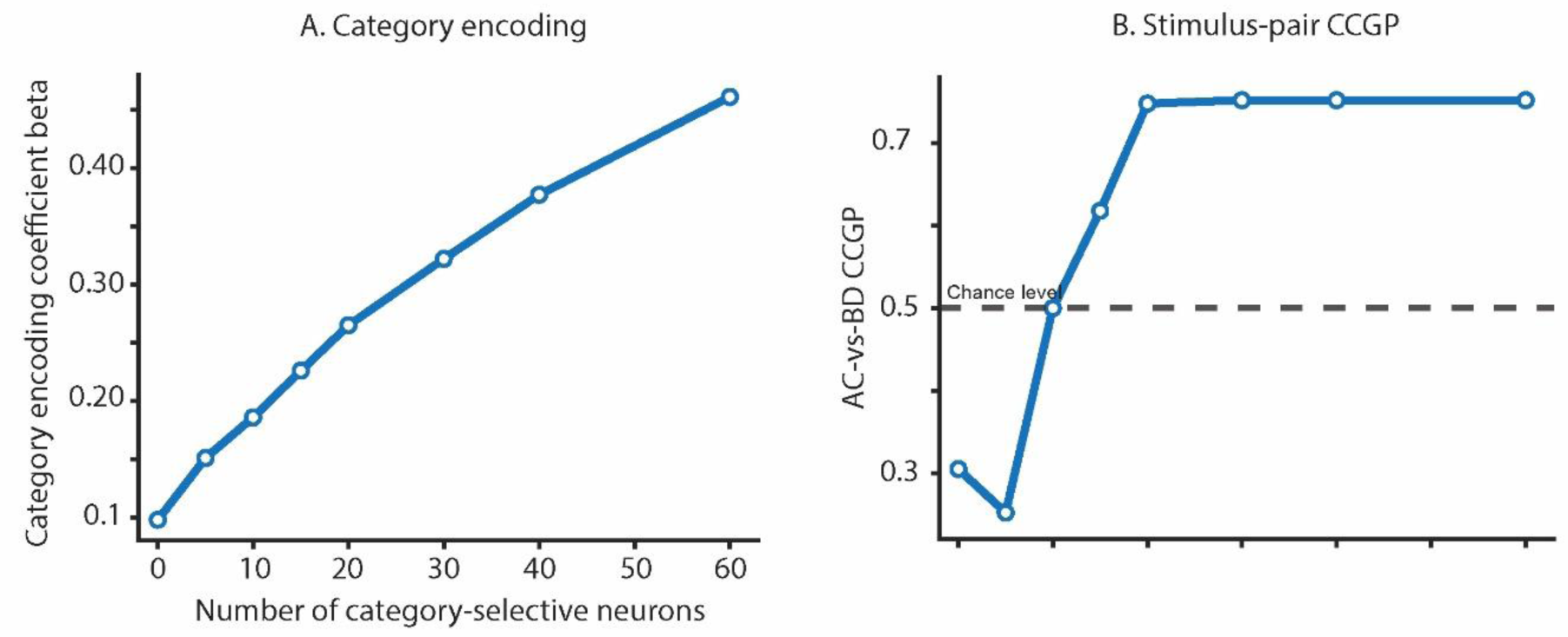
Category encoding and stimulus-pair cross-condition generalization as a function of category-selective neuron number. **(A)** Category encoding coefficient beta across increasing numbers of category-selective neurons. **(B)** AC-vs-BD stimulus-pair CCGP across increasing numbers of category-selective neurons. The dashed line indicates chance-level performance (CCGP = 0.5).

In parallel, AC-vs-BD stimulus-pair CCGP increased with increasing category-selective neuron number (Figure 1B). CCGP values increased from approximately 0.25–0.30 at low category-neuron numbers to approximately 0.75 with higher numbers of category-selective neurons. CCGP reached a plateau at approximately 20 category-selective neurons, while category encoding coefficient continued to increase with additional category-selective neurons.

### 3.3. Functional classification identifies category-like and identity-like hippocampal neurons

Functional classification of the hippocampal population revealed neurons with distinct task-related response profiles. Of the recorded neurons, 93 were classified as stimulus-related, 56 as context-related, 13 as response-related, 13 as reward-related, and 319 as non-selective. Within the stimulus-related population, 13 neurons exhibited category-like response profiles, whereas 80 exhibited identity-like response profiles.

### 3.4. Category-like and identity-like neuron manipulations differentially affect stimulus encoding and CCGP

To examine how the composition of hippocampal populations influences stimulus-pair CCGP and encoding strength, we progressively increased the number of category-like or identity-like neurons while maintaining the contribution of the remaining neuronal populations constant (Figure 2).

**Figure 2.**
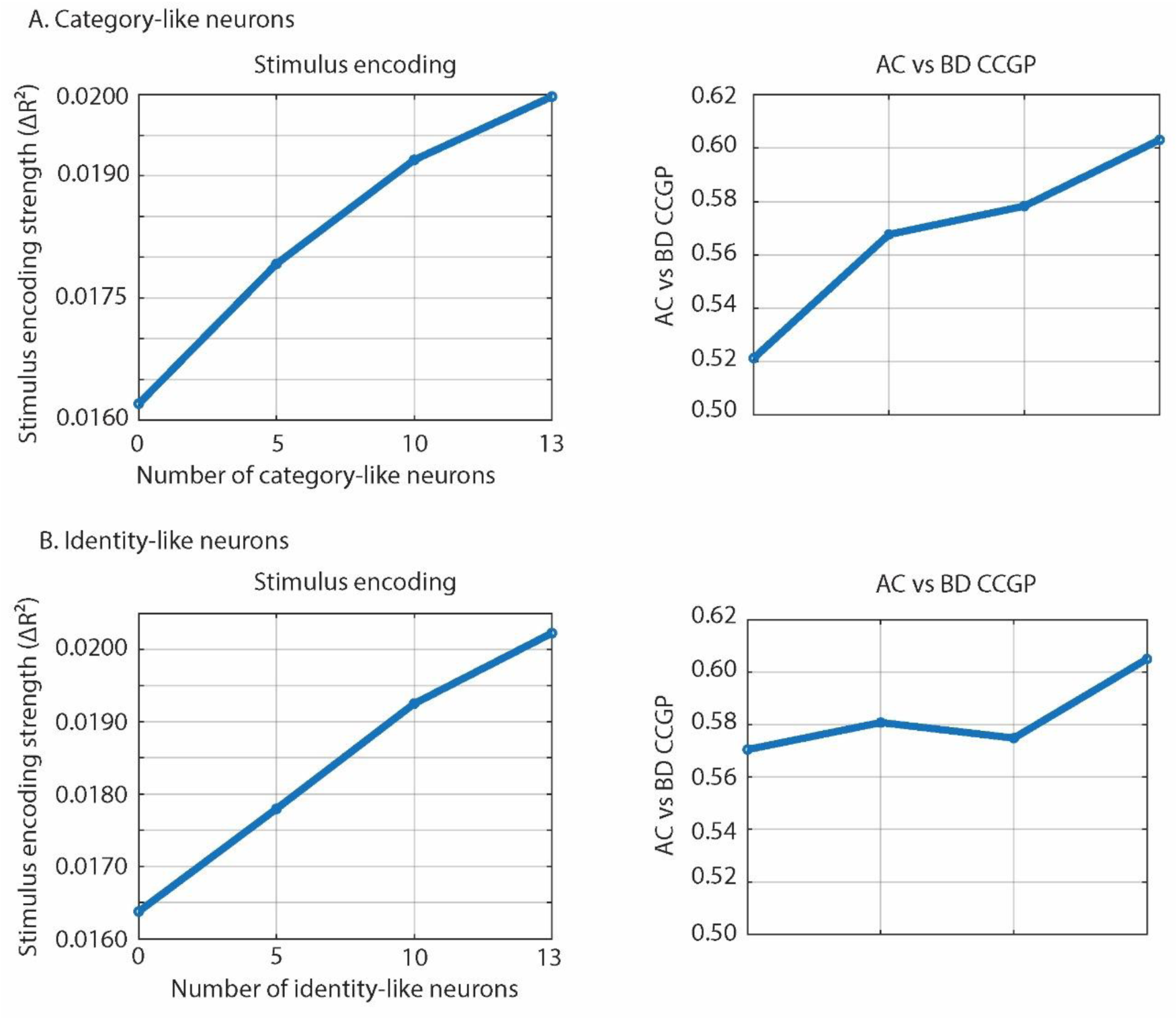
Stimulus encoding strength and AC vs BD CCGP across category-like and identity-like neuron manipulations. **(A)** Stimulus encoding strength (ΔR²) and AC vs BD CCGP values obtained by progressively increasing the number of category-like neurons in the population. **(B)** Stimulus encoding strength (ΔR²) and AC vs BD CCGP values obtained by progressively increasing the number of identity-like neurons in the population.

In the first manipulation, the number of category-like neurons was increased from 0 to 13 while identity-like neurons were fixed at 13. Stimulus encoding strength increased progressively from **0.0162** in the baseline population to **0.0199** when 13 category-like neurons were included. In parallel, AC vs BD CCGP increased from **0.521** to **0.605** across the same population configurations.

In the second manipulation, the number of identity-like neurons was increased from 0 to 13 while category-like neurons were fixed at 13. Stimulus encoding strength similarly increased from 0.0163 to 0.0202. However, AC vs BD CCGP showed a smaller increase, changing from 0.570 in the baseline condition to 0.605 with 13 additional identity-like neurons.

### 3.5. Neuronal composition alters AC–BD population geometry

Changes in neuronal composition were accompanied by distinct changes in the low-dimensional organization of the eight stimulus–context conditions (Figure 3). When category-like neurons were progressively added to a population containing 13 identity-like neurons, AC– BD separation in PC space increased across each successive configuration. The separation increased from approximately 2.54 with no category-like neurons to 3.02 with five category-like neurons, further increased to 3.41 with ten neurons, and reached 3.65 when 13 category-like neurons were included.

**Figure 3.**
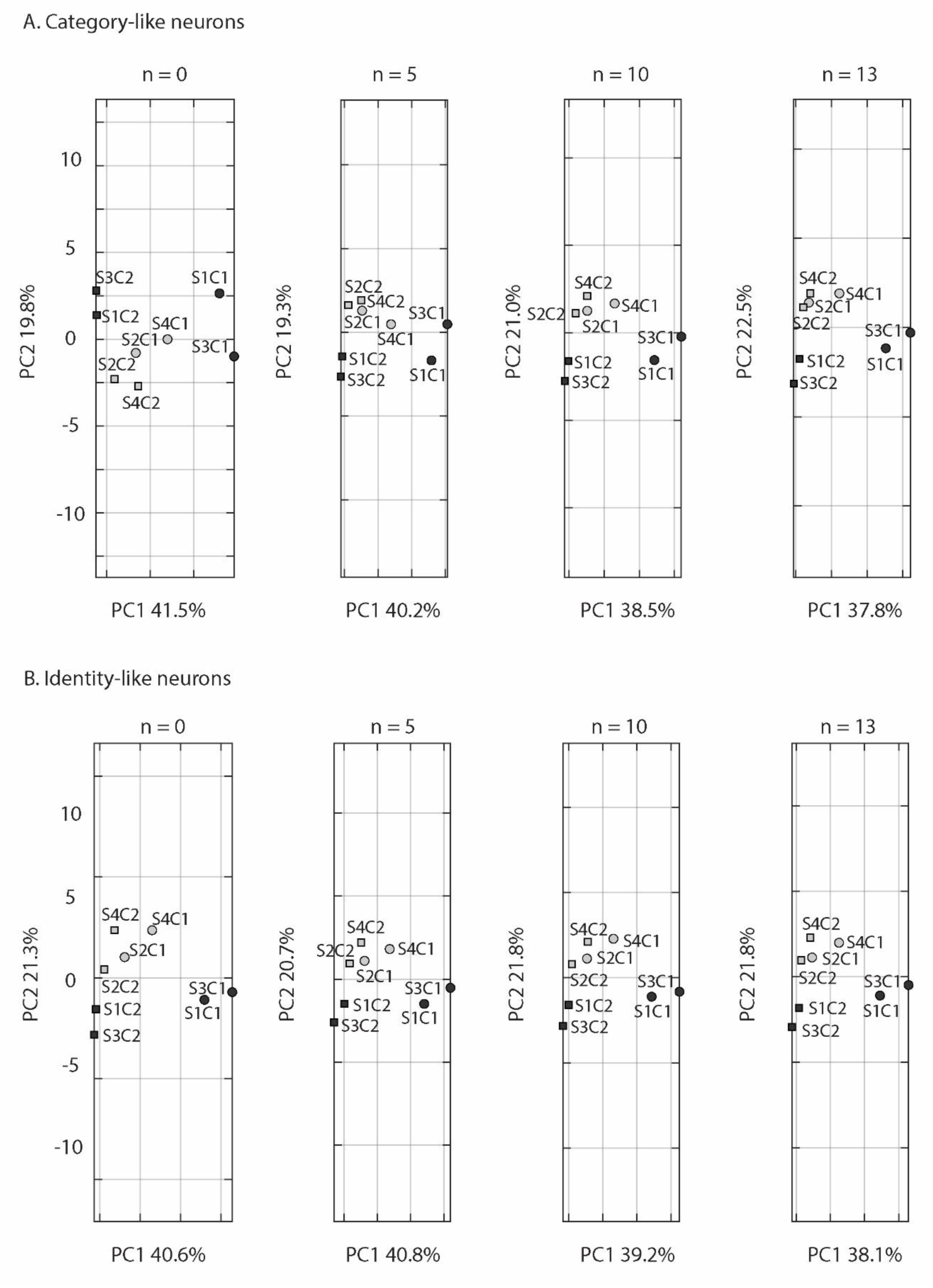
PCA projections of stimulus–context conditions across and PC-space distances. **(A)** Projection of the eight stimulus–context conditions onto the first two principal components as the number of category-like neurons was increased from 0 to 13, with 13 identity-like neurons retained in all populations. **(B)** Corresponding projections as the number of identity-like neurons was increased from 0 to 13, with 13 category-like neurons retained in all populations. In each row, n denotes the number of neurons of the manipulated class included in the pseudopopulation. Each point represents one of the eight stimulus–context conditions (S1C1–S4C2). Percentages along the axes indicate the variance explained by PC1 and PC2 for each population configuration.

The individual condition projections showed a progressive reorganization across the category-like neuron manipulation. In the absence of category-like neurons, the eight conditions were relatively close in PC space, with substantial overlap and limited separation between the S1/S3 and S2/S4 stimulus groupings. As category-like neurons were added, conditions belonging to the same stimulus grouping became more similarly positioned within each context, while the S1/S3 conditions became progressively separated from the S2/S4 conditions. This organization became increasingly apparent with the addition of 5, 10, and 13 category-like neurons. Stimuli belonging to the same category (S1/S3 and S2/S4) showed progressively similar spatial organization across contexts. At the same time, the two contexts became spatially segregated in the projection: in the 13-neuron configuration, the four C2 conditions occupied the left side of the PC1–PC2 space, whereas the C1 conditions were positioned further to the right. Thus, the stepwise addition of category-like neurons was accompanied by a progressive restructuring of both stimulus-pair and context-related organization in the PCA projection (Figure 3A).

A markedly different progression was observed when identity-like neurons were added while 13 category-like neurons were retained. Importantly, even in the absence of identity-like neurons, the eight conditions already displayed the structured arrangement observed after adding 13 category-like neurons in the preceding manipulation. The S1/S3 and S2/S4 groupings were separated, corresponding conditions showed similar relative arrangements within contexts, and the C1 and C2 conditions occupied distinct regions of the projected space. Adding 5 and subsequently 10 identity-like neurons produced comparatively small changes in this organization, without an evident progressive increase in the parallel arrangement of the corresponding stimulus groupings. With the final increase from 10 to 13 identity-like neurons, two conditions became nearly overlapping in the projection rather than becoming further separated (Figure 3B).

AC–BD separation varied only modestly across the identity-like neuron manipulation, from approximately 3.49 with no identity-like neurons to 3.31 with five neurons, 3.51 with ten neurons, and 3.61 when 13 identity-like neurons were included.

### 3.6. Category-axis alignment and strength show different relationships with CCGP

To determine which aspect of population geometry was most closely associated with stimulus-pair generalization, we examined the relationship between CCGP and two complementary geometric properties: category-axis alignment across contexts and category-axis strength. Across pseudopopulation realizations, CCGP showed a strong positive correlation with category-axis alignment (r = 0. 775, p < 0.001; Figure 4A), such that populations with more similarly oriented AC–BD axes across the two contexts exhibited higher cross-condition generalization. Category-axis strength was also positively correlated with CCGP, but the relationship was substantially weaker (r = 0. 370, p < 0.001; Figure 4B). Thus, both the consistency and magnitude of category separation covaried with CCGP at the bivariate level, with a considerably stronger association for cross-context axis alignment.

**Figure 4.**
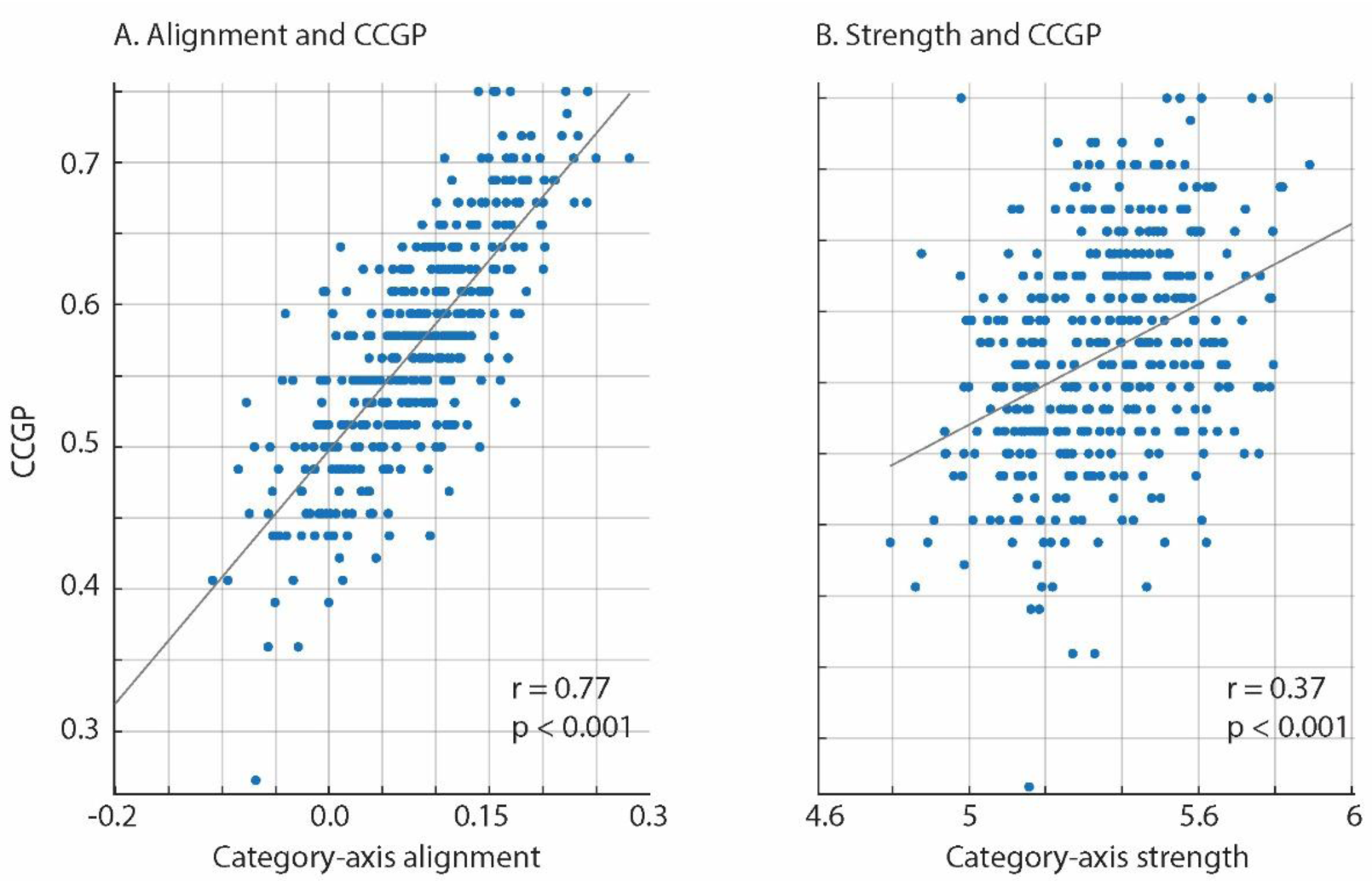
Relationships between category-axis geometry and CCGP. (A) Scatter plot of category-axis alignment against CCGP. (B) Scatter plot of category-axis strength against CCGP. Each point represents one pseudopopulation realization. Gray lines indicate linear regression fits. Pearson correlation coefficients (*r*) and corresponding *p*-values are displayed within each panel.

Partial-correlation analyses further examined the relationship of each geometric measure with CCGP while controlling for the other measure and neuronal composition. Category-axis alignment remained strongly associated with CCGP after controlling for category-axis strength and the numbers of identity-like and category-like neurons (partial r = 0.731, FDR-corrected p < 0.001). Category-axis strength also showed a statistically significant, though considerably weaker, partial association with CCGP (partial r = 0.11, FDR-corrected p = 0.023). These results show that, although both geometric properties were independently related to CCGP, the association was substantially stronger for category-axis alignment than for category-axis strength.

The robustness of these findings was further evaluated using two complementary sensitivity analyses. A neuronal-composition-level analysis assessed whether the zero-order associations were robust to resampling-level dependence, while a leave-one-category-neuron-out analysis assessed whether the differential partial associations of alignment and strength with CCGP were disproportionately influenced by any single category-like neuron. Results from both sensitivity analyses are reported in the Supplementary Results.

### 3.7. Mediation analysis of category-like neuron number, category-axis alignment, and CCGP

To determine whether the relationship between category-like neuron number and CCGP was mediated by changes in population geometry, we examined category-axis alignment as a mediator of this relationship. The mediation analysis showed that category-like neuron number was positively associated with category-axis alignment (path *a* = 0. 0037747, *p* < 0.001), such that populations containing more category-like neurons showed greater alignment of the category axis across contexts. Category-axis alignment was, in turn, positively associated with CCGP (path *b* = 0. 84638, *p* < 0.001). The total effect of category-like neuron number on CCGP was also significant (*c* = 0. 0033791, *p* < 0.001), corresponding to the increase in CCGP observed with increasing numbers of category-like neurons.

When category-axis alignment was included in the relationship, the remaining direct effect of category-like neuron number on CCGP was substantially reduced and was no longer significant (*c′* = 0. 00018, *p* = 0. 78647). The resulting indirect effect of category-like neuron number on CCGP through category-axis alignment was 0. 00319. Bootstrap analysis with 5,000 resamples produced a 95% confidence interval of [0. 00185, 0. 00448], which did not include zero (Figure 5). Thus, the increase in CCGP associated with adding category-like neurons was statistically accounted for by the accompanying increase in category-axis alignment. These results indicate that category-like neurons are associated with improved cross-context generalization primarily because their addition is accompanied by a more consistently aligned categorical geometry across contexts, rather than simply by an increase in the strength of category-related separation.

**Figure 5.**
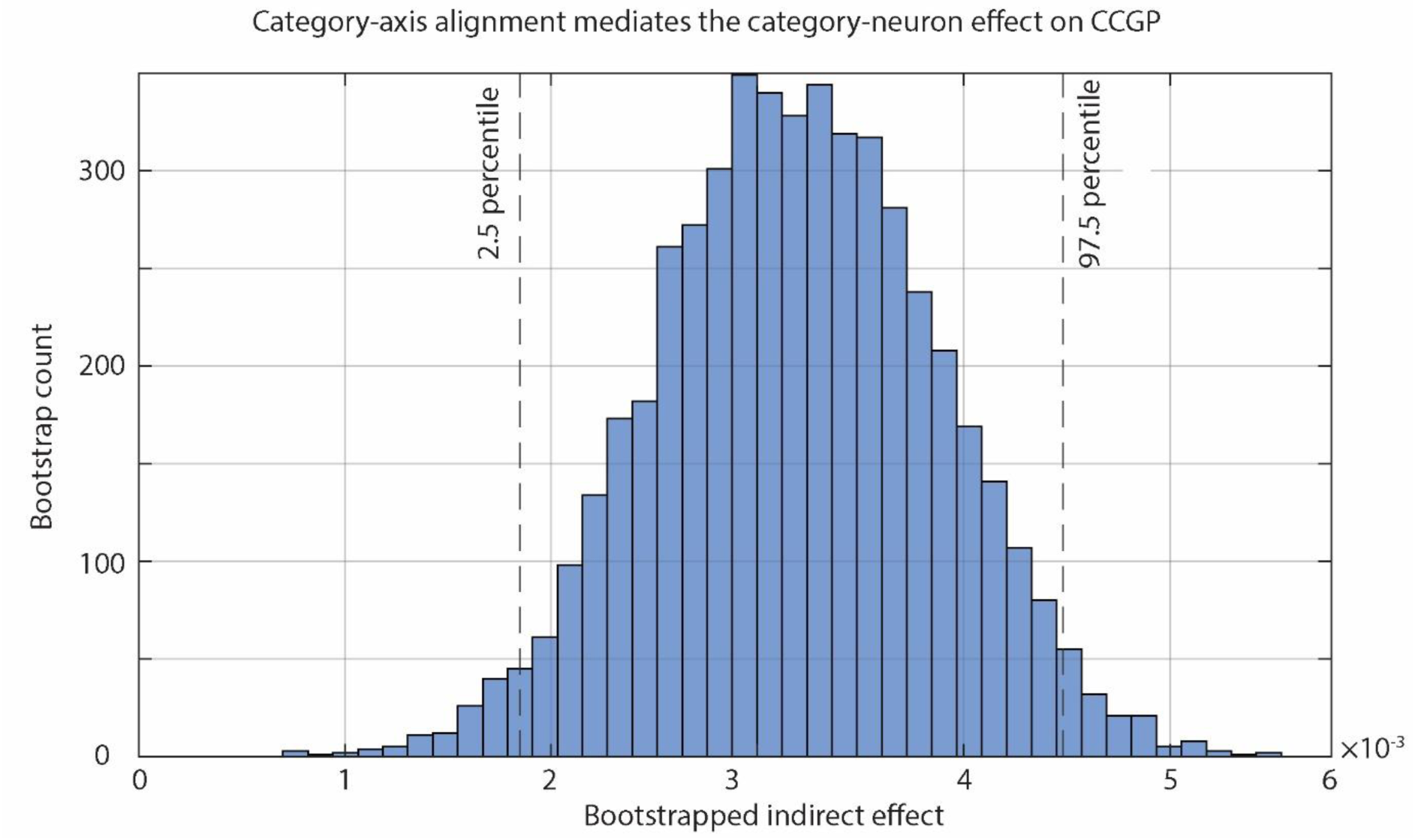
Bootstrap mediation analysis of category-axis alignment. Distribution of bootstrapped indirect effects for the pathway from category-like neuron number to CCGP through category-axis alignment. Dashed vertical lines indicate the 2.5th and 97.5th percentiles of the bootstrap distribution, corresponding to the bounds of the 95% bootstrap confidence interval.

## 4. Discussion

In this study, we investigated how neuronal population composition shapes the emergence of abstract representations that support generalization across contexts. Using a computational toy model and human hippocampal single-unit recordings, we found that increasing task-relevant information alone was not sufficient to improve abstraction. Instead, the effect depended strongly on the type of neurons contributing to the population. In the hippocampal data, adding category-like and identity-like neurons produced comparable increases in stimulus encoding, but category-like neurons resulted in a substantially greater improvement in CCGP. We further found that cross-context category-axis alignment was strongly associated with CCGP after accounting for category-axis strength and neuronal composition, whereas category-axis strength showed a substantially weaker independent relationship with CCGP. Finally, mediation analysis showed that increasing the number of category-like neurons increased category-axis alignment across contexts, which in turn was associated with higher CCGP.

### 4.1. Increasing task-related information is not sufficient for abstraction; neuronal selectivity matters

In the toy model, increasing the number of stimulus-and response-selective neurons strengthened stimulus and response encoding and increased the overall variance explained by the encoding model (R²), while AC-vs-BD CCGP remained at chance level (CCGP = 0.5). In contrast, introducing category-selective neurons increased CCGP above chance (CCGP = 0.619), together with further increases in task-related encoding. The subsequent category-neuron sweep further dissociated encoding strength from generalization: category encoding continued to increase as more category-selective neurons were added, whereas CCGP reached a plateau after approximately 20 category-selective neurons. This dissociation was reinforced by the real hippocampal data. Adding either category-like or identity-like neurons produced comparable increases in stimulus encoding, reaching the same encoding strength when 13 neurons of either type were included. However, category-like neurons produced a substantially greater increase in CCGP than identity-like neurons, with an approximately 2.5-fold greater improvement. The smaller increase in CCGP produced by identity-like neurons may reflect the fact that these neurons were not necessarily purely identity-selective. As expected in biological neural populations, neuronal selectivity was not strictly discrete, and neurons were classified as identity-like when their response profiles were better explained by stimulus identity than by category structure. Thus, some identity-like neurons may still have carried category-related information that contributed modestly to cross-context generalization. These results demonstrate that increases in task-related information do not necessarily translate into improved abstraction. Instead, the functional composition of the population determines whether additional information contributes to cross-context generalization.

This finding extends previous frameworks of abstract representation by suggesting a cellular mechanism through which population-level abstract geometries may emerge. Previous studies have shown that abstraction is associated with representations that preserve task-relevant relationships across changing conditions rather than with increased decodability alone [4, 5]. Our results suggest that such geometries are not determined solely by the amount of information represented in a population, but also by the selectivity structure of the neurons forming that population. Neurons whose activity aligns with latent categorical relationships can therefore disproportionately shape generalization, whereas neurons encoding stimulus-specific information may increase representational strength without producing equivalent improvements in abstraction.

Previous studies have shown that abstract representations can emerge through learning and inference, suggesting that experience can reshape neural population geometry to support flexible generalization [4, 5]. Our findings suggest that this reorganization may not involve all neurons equally, but may instead depend on the selective contribution of specific functional subpopulations. Specifically, because category-like neurons contributed disproportionately to CCGP despite producing similar increases in stimulus encoding as identity-like neurons, learning may promote abstraction by preferentially modifying or recruiting neurons whose activity is aligned with the latent structure of the task. This specialization could arise through two non-mutually exclusive mechanisms: learning may recruit pre-existing neurons with suitable representational properties, or experience-dependent plasticity may reshape neuronal tuning to generate category-like response profiles. Although our results cannot distinguish between these possibilities, they suggest that changes in neuronal selectivity and population composition may provide an intermediate link between learning-induced plasticity and the emergence of abstract representations.

A further observation from the toy model was that category encoding continued to increase as more category-selective neurons were added, whereas CCGP reached a plateau after approximately 20 neurons. This pattern raises the possibility that abstraction may require a sufficient, rather than continuously increasing, level of category-related information. However, this interpretation remains tentative because the plateau was observed only in the toy model. The limited number of category-like neurons available in the empirical dataset precluded an equivalent extended sweep in the human data, preventing us from determining whether a similar saturation would emerge in biological populations. Importantly, this constraint is specific to testing the saturation effect predicted by the toy model and does not apply to the other empirical analyses.

### 4.2. A relatively small category-like subpopulation can disproportionately reshape population representation

In the real hippocampal data, adding a relatively small number of category-like neurons (13 neurons) produced a substantially larger reorganization of population geometry than adding a much larger population of identity-like neurons (∼80 neurons). Specifically, category-like neurons progressively increased AC–BD separation in PCA space, whereas identity-like neurons produced only modest changes despite contributing comparable increases in stimulus encoding. This finding suggests that the functional influence of a neuronal subpopulation is not necessarily proportional to its numerical prevalence within a population.

This observation is consistent with previous studies in the human medial temporal lobe (MTL) demonstrating that sparse neuronal populations can carry highly selective and invariant representations of complex information. Human MTL neurons have been shown to respond selectively across different exemplars of the same individual, object, or concept, suggesting that relatively small neuronal populations can encode high-level invariant representations [13, 14]. However, previous work has primarily focused on the contribution of sparse neurons to what information is represented. Our findings extend this perspective by suggesting that sparse functional subpopulations may also influence how information is organized at the population level. Specifically, a minority population of neurons with selectivity aligned with latent task structure may disproportionately shape the geometry of the neural representation, thereby promoting abstract cross-context generalization.

Beyond their role in supporting abstraction, different neuronal subpopulations may contribute differently to the balance between stimulus-specific representation and generalization. Identity-like neurons may preferentially preserve information about individual stimuli, whereas category-like neurons may promote representations of shared latent structure. Therefore, variation in the relative composition of these neuronal populations could provide a mechanism by which hippocampal representations are tuned toward different points along the specificity– generalization continuum. This interpretation aligns with computational studies showing that representational geometry determines the balance between supporting abstraction and generalization while preserving flexible information coding [6, 9]. For example, Boyle et al. [9] demonstrated that hippocampal representations can adopt different geometries that satisfy distinct computational demands, with some geometries favoring detailed memory representations and others supporting broader generalization. Our findings extend this framework by suggesting that the emergence of such geometries may be partly shaped by the composition of the underlying neuronal population. Specifically, the relative contribution of neurons with different selectivity profiles may influence the organization of population activity and thereby determine the computational regime of the representation.

The ability of relatively small neuronal subpopulations to reshape population-level representations raises the possibility that differences in cognitive performance may partly arise from variation in the composition or functional organization of neural populations. Specifically, individuals may differ in the extent to which neurons with selectivity aligned to latent task structure are recruited or strengthened during learning, leading to differences in the geometry of the resulting representations and, consequently, in the ability to generalize beyond individual experiences. This possibility is consistent with previous studies showing that learning can shape the emergence of abstract conceptual representations and reorganize neural population structure [15, 16]. In this framework, more efficient formation of category-aligned representations could favor abstract inference and flexible knowledge transfer, whereas stronger preservation of stimulus-specific representations may favor detailed memory for individual experiences at the expense of broader generalization. Importantly, this hypothesis does not imply that individuals with better memory or inference abilities simply have more category-like neurons. Rather, it suggests that individual differences may arise from how neuronal populations are organized and balanced along the specificity–generalization continuum. Future longitudinal studies combining single-neuron population dynamics with behavioral measures of learning and inference will be needed to determine whether differences in the recruitment and stabilization of functionally specialized subpopulations contribute to variability in cognitive flexibility.

### 4.3. Cross-context alignment, rather than representational strength, is the geometric property most closely associated with abstraction

A central finding of our study is that the ability of a population representation to support abstraction was more closely related to the preservation of category structure across contexts than to the magnitude of category-axis separation within individual contexts. Although both category-axis strength and alignment were positively associated with CCGP, their contributions differed substantially. Category-axis alignment showed a strong relationship with cross-context generalization that remained robust after controlling for category-axis strength and neuronal composition, whereas the association between category-axis strength and CCGP was considerably weaker (partial r = 0.11 compared with 0.73 for alignment). These results suggest that strong category representations are not necessarily abstract representations; rather, abstraction is supported when the geometric relationships that define a category are preserved across changes in context.

This distinction reflects a fundamental property of abstract neural representations. A population may contain highly separable representations within each context, yet fail to generalize if the dimensions that separate categories are reorganized across contexts. In contrast, when the relevant representational axes remain aligned, downstream neural systems can apply similar readout mechanisms across different situations without requiring a new decoding strategy for each context. Thus, abstraction may emerge not simply from increasing the magnitude of category-related information, but from maintaining a stable geometric organization that preserves task-relevant relationships across experiences.

This interpretation is consistent with computational frameworks showing that representational geometry determines whether neural populations support flexible generalization while maintaining rich information content [5,12]. Rather than requiring identical neural activity patterns across contexts, abstract representations can preserve relationships among representations through shared geometric structure. Our findings extend this framework by suggesting that neuronal population composition may influence abstraction by shaping the alignment of these representational structures. In this view, category-like neurons do not promote abstraction merely by increasing category-related signal; instead, their contribution may arise because they help establish a population geometry in which category relationships remain stable across contexts.

The strong relationship between cross-context alignment and CCGP also provides a possible interpretation of the plateau observed in the toy model. In that analysis, category encoding continued to increase as more category-selective neurons were added, whereas CCGP showed little further improvement beyond approximately 20 category neurons. These results raise the possibility that learning may prioritize the organization of task-relevant information within representational space rather than continuously increasing the magnitude of neural responses. In this framework, learning may not improve abstraction by simply strengthening category-related activity, but by refining the geometric arrangement of representations so that relevant relationships remain stable across changing contexts. Such a mechanism would provide an efficient solution for generalization, allowing existing neural readouts to be transferred across experiences without requiring progressively stronger encoding signals. This hypothesis can be directly tested in future studies by tracking changes in population geometry across different stages of learning and determining whether improvements in abstraction are better explained by increases in representational alignment than by changes in encoding strength.

### 4.4. Category-like neurons support abstraction primarily through geometric alignment

The mediation analysis provides a potential mechanistic link between neuronal selectivity and population-level abstraction. Although the presence of category-like neurons significantly predicted increases in CCGP, this relationship was largely explained by changes in cross-context alignment. Specifically, category-like neuron composition significantly predicted alignment, and alignment strongly predicted CCGP, whereas the direct effect of category-like neurons on CCGP was no longer significant after accounting for alignment. These results suggest that category-like neurons may not promote abstraction simply by adding category-related information to the population, but rather by shaping the geometry through which this information is organized across contexts.

This finding provides a possible bridge between single-neuron selectivity and population-level computational properties. Previous studies have separately emphasized the importance of neuronal selectivity in representing meaningful variables [13] and the role of population geometry in supporting abstraction and generalization [4–6]. Our results suggest a potential mechanism linking these two levels: neuronal selectivity may influence computation because it determines the geometric structure of the population representation, which in turn governs the ability of the system to generalize across contexts. Importantly, given that the mediation analysis is based on resampled population manipulations rather than direct experimental intervention, these results should be interpreted as evidence consistent with an intermediate role for geometry rather than proof of causality.

The mediation analysis results complement our earlier interpretation regarding the potential role of neuronal population composition in individual differences in abstraction and generalization. We previously proposed that individuals may differ in how effectively category-aligned neuronal responses are recruited or strengthened during learning, potentially shifting hippocampal representations toward different points along the specificity– generalization continuum. The present results suggest a possible population-level mechanism through which such differences could influence cognitive performance: variation in the formation or recruitment of functionally appropriate neuronal subpopulations may lead to differences in how effectively task-relevant representations become geometrically aligned across contexts. Thus, individual variability may ultimately depend not only on which neuronal responses emerge during learning, but also on how effectively these responses reorganize population geometry to support transfer [15, 17]. This hypothesis remains to be tested directly and will require longitudinal studies tracking neuronal selectivity, geometric alignment, and behavioral generalization throughout learning.

### 4.5. Population geometry as a computational principle for improving artificial generalization

Beyond biological insight, our findings suggest potential principles for improving generalization in artificial neural networks. Modern artificial systems are typically optimized by maximizing task performance through feature learning and loss minimization [18], with improvements often driven by increasing model capacity or the amount of task-relevant information encoded within network representations. However, our results suggest that information quantity alone may not be sufficient for robust generalization across changing contexts [4]. In the toy model, increasing stimulus-and response-related information substantially improved encoding performance but failed to enhance cross-context generalization, whereas introducing category-selective representations increased CCGP. These findings suggest that the organization of task-relevant information within representational space, rather than its magnitude alone, may be a critical determinant of transfer [4, 19].

This perspective raises the possibility that artificial networks could achieve more robust out-of-context generalization by incorporating objectives that explicitly promote cross-context representational alignment. Instead of optimizing only for accurate task outputs, future learning algorithms could combine task objectives with geometry-aware constraints that encourage shared latent structures to remain aligned across different contexts [18, 20]. Such approaches could potentially produce representations that maintain transferability without requiring separate adaptation for each new environment.

Our results further suggest that increasing representational strength or network capacity may provide diminishing returns when the geometry of learned representations is not appropriately organized. The category-neuron sweep demonstrated that category-related encoding could continue increasing even after CCGP reached a plateau, indicating that additional information does not necessarily translate into improved generalization. A similar principle may apply to artificial systems: larger or more expressive representations may not automatically yield better transfer unless the learned geometry preserves task-relevant relationships across contexts [4, 19].

Finally, the finding that a relatively small category-like neuronal subpopulation could disproportionately reshape population geometry suggests a possible role for functional specialization in artificial networks. Effective generalization may not require all units to develop invariant representations; instead, a specialized subset of units may be sufficient to organize population activity into a geometry that supports transfer. This raises the possibility that future architectures could benefit from heterogeneous representations in which some units preserve stimulus-specific information, whereas others encode higher-order relational structure [21]. Such a balance between specificity-preserving and abstraction-supporting components may provide a computational solution for maintaining detailed representations while enabling flexible generalization.

### 4.6. Limitation

A further consideration concerns the statistical structure of the pseudopopulation analyses. Because pseudopopulation realizations were repeatedly constructed from a finite pool of recorded neurons, individual realizations should not be interpreted as fully independent biological samples. This is an inherent constraint of human single-neuron datasets, which provide exceptionally detailed access to neuronal activity but are rare and necessarily limited in the number of available neurons, particularly for experimental paradigms suitable for studying abstract generalization. We therefore evaluated the robustness of the geometry–CCGP findings using complementary composition-level and leave-one-neuron-out sensitivity analyses. These analyses supported the robustness of the observed associations but do not create additional independent biological observations. Accordingly, the partial-correlation and mediation analyses should be interpreted as evidence for statistical relationships within the sampled neuronal population rather than as definitive estimates of population-level effects. In particular, the mediation analysis is consistent with a potential indirect relationship linking category-like neuronal composition, cross-context alignment, and generalization, but should not be interpreted as establishing a causal mediation mechanism.

A separate limitation applies specifically to testing the saturation effect observed in the toy model. In the simulations, the availability of an unrestricted synthetic neuronal population allowed us to systematically increase the number of category-like neurons over a broad range, revealing that CCGP tended to plateau despite continued increases in category-related encoding. The empirical dataset contained 13 neurons classified as category-like, which limited the range over which an analogous neuronal-composition sweep could be performed in the human data. Consequently, the real-data analyses cannot determine whether further increases in the number of category-like neurons would eventually produce a similar saturation of generalization. Importantly, this limitation is specific to testing the predicted saturation effect and does not constrain the other empirical analyses reported here, which were performed using the complete available neuronal dataset and directly characterize the observed relationships among neuronal selectivity, representational geometry, and generalization within that dataset.

## 5. Conclusion

In conclusion, our findings identify how neuronal populations may generate abstract representations that support generalization across changing contexts. By combining controlled manipulations in a computational model with analyses of human hippocampal population activity, we show that abstraction does not depend simply on having more task-related information in a neural population. Instead, it depends on which neurons contribute this information and how their activity organizes the population representation. Neurons with selectivity that matches the underlying structure of the task can have a strong influence on the emergence of generalizable representations, not only by increasing category-related information but also by promoting a consistent geometric organization across contexts. These findings suggest that successful generalization relies on preserving important relationships between experiences rather than simply increasing the strength of neural representations. More broadly, our study provides a link between single-neuron selectivity and population-level computation, suggesting how learning-related changes in neuronal populations may shape the emergence of flexible and abstract representations. This principle may also provide useful guidance for developing artificial systems that require robust generalization beyond the conditions in which they were trained.

## Data availability

The human hippocampal single-unit dataset analyzed in this study was obtained from the publicly available dataset accompanying Courellis et al., *Nature* (2024) (https://doi.org/10.1038/s41586-024-07799-x; https://doi.org/10.17605/OSF.IO/QPT8F). No new experimental data were collected for this study.

## Code availability

All custom code used in this study is publicly available on GitHub at (https://github.com/ArminH69/Neuronal-selectivity-for-CCGP). The repository contains the Python code for the computational toy-model analyses and the MATLAB scripts used for the re-analysis of human hippocampal single-unit recordings.

## Acknowledgments

We thank Professor Alexandre Hyafil for his valuable guidance and discussions regarding the analyses.

## Author Contributions

A.H.M.T. conceived the study, developed the methodology, performed the analyses, interpreted the results, and wrote the manuscript. N.D. contributed to the computational toy-model analyses, validation of results, and manuscript review and editing.

## Supplementary Results

### S1. Sensitivity analysis at the neuronal-composition level

To assess whether the observed zero-order relationships between population geometry and CCGP depended on treating individual pseudopopulation resampling iterations as separate observations, we repeated the correlation analyses after aggregating the resampling iterations within each neuronal composition. Each composition was defined by the combination of category-like and identity-like neuron numbers, resulting in four composition-level observations. For each composition, category-axis alignment, category-axis strength, and Category CCGP were averaged across the 100 resampling iterations.

The positive association between category-axis alignment and CCGP was preserved after aggregation at the neuronal-composition level (r = 0.991, p = 0.009). Category-axis strength was also strongly positively correlated with CCGP at the composition level (r = 0.995, p = 0.005). Thus, the positive zero-order relationships observed in the resampling-level analysis were not dependent on treating the individual resampling iterations as separate observations.

Notably, this sensitivity analysis was designed specifically to evaluate the robustness of the zero-order associations to resampling-level dependence and was not intended to distinguish the independent contributions of alignment and strength. Only four neuronal compositions were available, and across these compositions, increasing the number of category-like neurons was accompanied by concurrent increases in category-axis alignment, category-axis strength, and CCGP. Consequently, the composition-level correlations primarily capture their shared variation across neuronal compositions and cannot determine whether alignment or strength is independently more strongly associated with CCGP. Their independent relationships with CCGP were instead evaluated in the primary partial-correlation analysis, which controlled for the alternative geometric measure and neuronal composition.

Overall, the composition-level analysis provides a sensitivity check showing that the positive geometry–CCGP associations, including the strong alignment–CCGP relationship, persist when repeated pseudopopulation realizations are collapsed into single composition-level observations. These results therefore support the robustness of the zero-order associations to the number of resampling iterations, while the relative independent relationships of alignment and strength with CCGP are addressed by the primary partial-correlation analysis.

### S2. Leave-one-category-neuron-out sensitivity analysis

To assess whether the stronger partial association between category-axis alignment and CCGP was disproportionately influenced by any individual category-like neuron, we performed a leave-one-category-neuron-out sensitivity analysis. Each of the 13 category-like neurons was removed in turn, and the pseudopopulation analysis was repeated using the remaining 12 category-like neurons. For each leave-one-out dataset, four neuronal compositions containing 0, 5, 10, or 12 category-like neurons were evaluated while the number of identity-like neurons was fixed at 13, with 100 pseudopopulation realizations generated for each composition.

The alignment–CCGP partial association was highly stable across all 13 leave-one-out analyses. The mean partial correlation was r = 0.724 (median = 0.721), with values ranging from 0.691 to 0.776 across the 13 analyses. In contrast, the corresponding partial association between category-axis strength and CCGP remained consistently small, with a mean partial correlation of r = 0.054 (median = 0.051) and a range from −0.006 to 0.144.

Accordingly, the difference between the alignment–CCGP and strength–CCGP partial correlations remained positive in every leave-one-out analysis. The mean difference in partial correlation coefficients was Δr = 0.670 (median = 0.680), with a range from 0.575 to 0.727, and the alignment–CCGP partial correlation exceeded the strength–CCGP partial correlation in 13/13 leave-one-out analyses (100%).

These results indicate that the substantially stronger independent association of category-axis alignment with CCGP was not driven by any single category-like neuron. Removal of individual category-like neurons produced only limited variation in the alignment–CCGP association, which remained strong in every analysis, whereas the strength–CCGP association remained weak. This leave-one-neuron-out analysis therefore provides an influence-based robustness check complementary to the neuronal-composition-level sensitivity analysis in S1. Whereas S1 assessed whether the zero-order geometry–CCGP associations persisted after removing resampling-level replication, the present analysis assessed whether the differential partial-correlation pattern was disproportionately dependent on individual category-like neurons.

